# Trans-synaptic dwelling of SARS-CoV-2 particles perturbs neural synapse organization and function

**DOI:** 10.1101/2022.09.13.507484

**Authors:** Emma Partiot, Aurélie Hirschler, Sophie Colomb, Willy Lutz, Tine Claeys, François Delalande, Maika S. Deffieu, Judith R.E. Roels, Joanna Bons, Domitille Callon, Laurent Andreoletti, Marc Labrousse, Frank M.J. Jacobs, Valérie Rigau, Benoit Charlot, Lennart Martens, Christine Carapito, Gowrishankar Ganesh, Raphael Gaudin

## Abstract

SARS-CoV-2 infection is associated with short- and long-term neurological and psychiatric complications, referred to as neuroCOVID. These symptoms are relatively heterogenous and fluctuating, hampering the discovery of molecular mechanisms underlying viro-induced brain perturbations. Here, we show that the human cerebral cortex poorly supports SARS-CoV-2 dissemination using *post-mortem* COVID-19 patient samples, *ex vivo* organotypic cultures of human brain explants and stem cell-derived cortical organoids. Despite restricted infection, the sole exposure of neural cells to SARS-CoV-2 particles is sufficient to induce significant perturbations on neural synapse organization associated to electrical activity dysfunction. Single-organoid proteomics revealed that exposure to SARS-CoV-2 is associated to trans-synaptic proteins upregulation and unveiled that incoming virions dwell at LPHN3/FLRT3-containing synapses. Our study provides new mechanistic insights on the origin of SARS-CoV-2-induced neurological disorders.

**One-Sentence Summary:** SARS-CoV-2 modulates neural plasticity and electrical activity as viral particles lodge at the trans-synaptic interface.

## Main Text

The coronavirus disease 2019 (COVID-19) results from the severe acute respiratory syndrome coronavirus 2 (SARS-CoV-2), which has caused an unprecedented worldwide pandemic. The most recognized organ to be impacted by the infection is the lower respiratory tract, but increasing number of studies indicate that the infection has also wide consequences on the central nervous system (CNS) (*1*). COVID-19 patients exhibit a large range of neurological, cognitive and psychiatric disorders (*2*–*4*), and 30-67% patients present neuropsychological impairments (*5*–*7*) as well as persistent abnormal electroencephalograms (*8*, *9*). Moreover, altered mental status has been widely reported as a long-term symptom of neuroCOVID (*10–15*). Modest neuroinflammation has been reported in patients with severe COVID-19 (*16*), which do not correlate with symptoms (or absence thereof). Thus, the relatively subtle neurological symptoms that have been described for the wide majority of patients with benign COVID-19 indicate that immunologically-independent neural perturbations must also exist. Here, we hypothesized that neural perturbations must also originate from other molecular and subcellular events that have not been uncovered yet because of the heterogeneity of the models and their limitations (*17*–*20*). Here, we propose a novel paradigm to explain viro-induced neurological disorders, where viral particles would interfere physically with trans-synaptic structures, leading to local electrical perturbations. These results provide an integrated explanation for most of the diverse results reported in the previous studies and open the way to the development of novel therapeutic strategies to treat neuroCOVID.

### SARS-CoV-2 infection of the human brain cortex

Our anatomopathological observations of the frontal and parietal lobes of human samples, obtained from patients who died from COVID-19 in the absence of neurological symptoms, did not allow us to observe astrogliosis, inflammatory infiltrates nor macroscopic abnormalities (**Fig. 1A-B** and **Suppl Fig. S1A-B**). As in most previous studies, we did not detect SARS-CoV-2 antigens by western blot, but we could identify trace amount of viral RNA in the temporal lobe, even in patients with negative blood viremia (**Suppl Fig. S1C**), indicating that some virus could reach and potentially persist into the brain as recently suggested (*21*).

**Figure 1.**
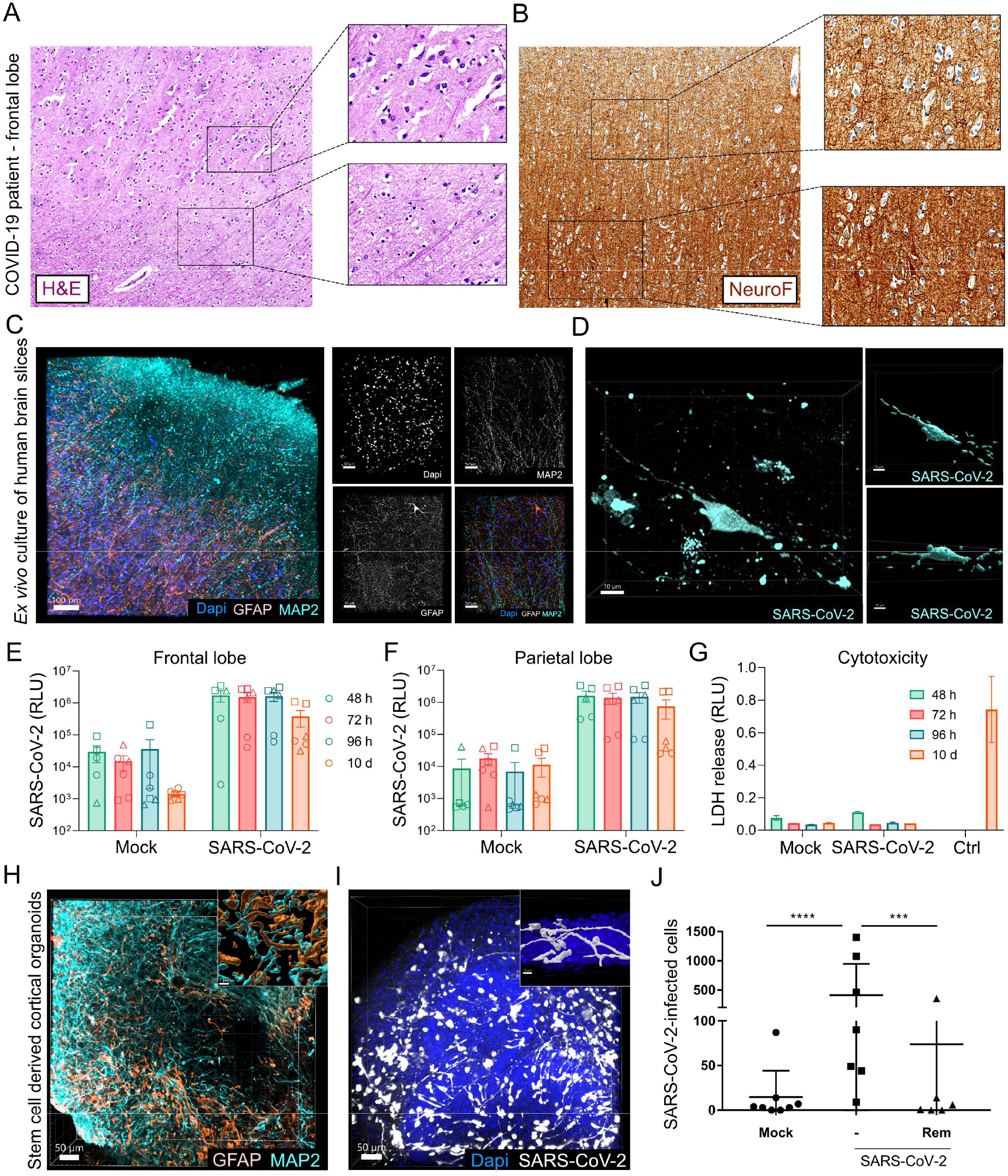
SARS-CoV-2 infection in the human cortex. (**A-B**) Hematoxylin and eosin staining (H&E, **A**) and immunohistochemistry of neurons (NeuroF, **B**) of samples obtained from the frontal lobe of a COVID-19 patient that did not present neurological symptoms. Images were acquired using a 20X objective. (**C-G**) Organotypic *ex vivo* culture of *post mortem* human brain slices mock-infected or infected with SARS-CoV-2 at 10^5^ pfu/well. (**C**) Three-dimensional snapshots of noninfected parietal lobe slices show the organization of mature neurons (MAP2) and astrocytes (GFAP). (**D**) Three-dimensional snapshots of a slice infected with a replication-competent SARS-CoV-2 mNeonGreen reporter virus. (**E-F**) Infection kinetics of frontal (**E**) and parietal (**F**) lobe slices upon infection with a replication-competent SARS-CoV-2 NanoLuciferase (NLuc) reporter virus. Each dot corresponds to a single slice and the symbols discriminate between the donors. (**G**) Cytotoxicity is measured for each condition from the parietal lobe using an LDH release luminescence assay. (**H**) Three-dimensional snapshots of cortical organoids stained for mature neurons (MAP2) and astrocytes (GFAP). (**I-J**) Cortical organoids were mock-infected or infected for 10 days with SARS-CoV-2 mNeonGreen at 10^5^ pfu/organoid in the presence or absence of 10 μM Remdesivir. The micrograph (**I**) shows a highly infected organoid (white staining) and nuclei (blue). The graph (**J**) corresponds to the number of infected cells from 7 individual organoids per condition. Two-tailed p value was < 0.001 (***) or < 0.0001 (****).

To better control the impact of SARS-CoV-2 exposure to the human brain, we developed an organotypic *ex vivo* culture of frontal and parietal cortex slices obtained from *post-mortem* brain resection of non-COVID patients (**Fig. 1C and Suppl Fig. S2**). Anatomopathological observations showed lower neuronal density after cortical slicing (**Suppl Fig. S2B**), likely due to the cutting procedure. Neural cell viability was mostly preserved beyond the edge of the slice and overall tissue architecture and cellular composition were maintained in the slices (**Fig. 1C** and **Suppl Fig. S2C-E**). The functional activity of cultured brain slices was readily monitored as they exhibited significant spontaneous electrical activity as measured by 3D microelectrode array (3D-MEA; **Suppl Fig. S2F-G**), further indicating that this model as retained functional properties. *Ex vivo* infection of parietal and frontal brain slices with SARS-CoV-2 reporter viruses (*22*) highlighted that a marginal number of cells could be infected, and the virus could not further propagate into the slice after initial virus inoculation (**Fig. 1D-F**). Moreover, the addition of SARS-CoV-2 to the slices did not result in toxicity nor major tissue disorganization (**Fig. 1G** and **Suppl Fig. S2E**). This data obtained in a physiologically relevant human model highlights that SARS-CoV-2 can infect neural cells to a very low extent, although they are unlikely to support the production of new infectious particles as viral replication kinetics reached a plateau after the initial inoculum. The low number of positive cells prevented us from robustly characterizing virus tropism in this model. Organotypic cultures have complex immune environment that is difficult to control for, and therefore, to gain further insights into the replication of SARS-CoV-2 in neural cells, we next use cortical organoids differentiated from stem cells, a 3D system that models an embryonic cortex, although devoid of microglia. Cerebral organoids exhibit neural progenitor cells, mature neurons, and astrocytes (**Fig. 1H** and **Suppl Fig. S3A**). Confocal image quantification showed the important heterogeneous SARS-CoV-2 permissiveness of this model, ranging from barely to highly infected status (**Fig. 1I-J** and **Suppl Fig. S3B**), reminiscent of the variability previously reported in cerebral organoid models (*17*). Moreover, the kinetics of SARS-CoV-2 replication in organoids was stalled (**Suppl Fig. S3C**), similar to our observations in brain slices (**Fig. 1E-F**). We confirmed that mature neurons were the most permissive cell type (**Suppl Fig. S3D**), as previously reported (*23, 24*), although SARS-CoV-2 cellular neurotropism is a complex question that is still under investigation (*25*, *26*). SARS-CoV-2 infection of cortical organoids does not lead to significant cytotoxicity, apoptosis, nor organoid growth defect (**Suppl Fig. S3E-G)**. Further characterization of cortical organoid infection showed replication-dependent expression of the viral protein N, while the organoids did not release new infectious particles (**Suppl Fig. S3H-I**). Together, our data indicates that the cortical regions of the brain represent a poorly permissive dissemination compartment for SARS-CoV-2. As such, we hypothesize that exposure of the brain to SARS-CoV-2 would cause local and transient perturbations, a scenario consistent with the pleiotropic and fluctuating neurological symptoms observed in most COVID-19 patients.

### SARS-CoV-2 perturbs neural synapse composition and activity

Although SARS-CoV-2 infection is relatively inefficient in the brain, the impact of virus exposure to the cortex remains to be mechanistically elucidated. To identify the molecular perturbations caused by SARS-CoV-2, we performed mass spectrometry-based differential proteomics on single cortical organoids infected in the presence/absence of virus (**Fig. 2A** and **Table S1**), allowing us to take into account the organoid-to-organoid variable permissiveness of SARS-CoV-2. Due to the heterogeneous nature of this approach, we could not reliably cluster infected and non-infected replicates together (**Suppl Fig. S4A**). Nevertheless, deeper analyses using dedicated bioinformatic tools allowed us to extract new relevant information. First, we compared the post-translational modifications (PTM) of the proteomes and found numerous proteins showing remarkable SARS-CoV-2-dependent PTM (**Suppl Fig. S4B-E** and **Table S2**). Moreover, we report the deamidation of the asparagine on position 29 of the viral N protein, a previously unknown modification for which no function has been described to our knowledge. The PTM from the proteome of cortical organoids, and how SARS-CoV-2 modulates them, represents a valuable dataset which meaning remains to be functionally investigated.

**Figure 2.**
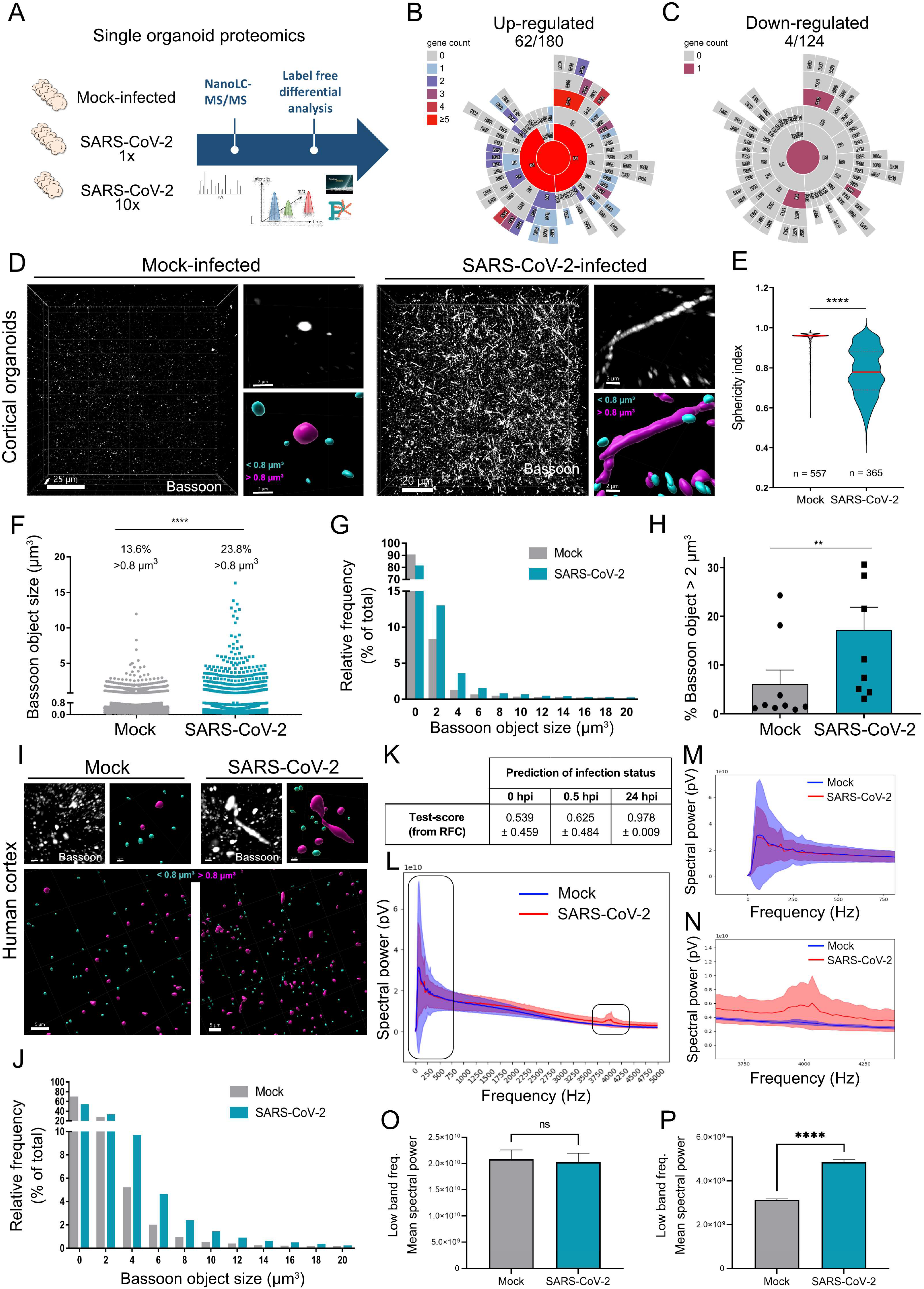
SARS-CoV-2 alters neural synapse morphology and function. (**A**) Cortical organoids were mock-infected or infected for 10 days with SARS-CoV-2 at 10^5^ pfu (1x) or 10^6^ pfu (10x) per organoid and processed for single-organoid differential proteomics comparing infected organoids to their non-infected counterparts using nanoLC-MS/MS. (**B-C**) The upregulated (**B**) and downregulated (**C**) proteins were analyzed using SynGo database and diagrams were built based on the synaptic distribution of the proteins belonging to the synaptosome. The color code corresponds to the number of synaptic proteins present in each sub-compartment. (**D**) Cerebral organoids were mock-infected or infected for 10 days with SARS-CoV-2 at 10^5^ pfu and stained for the presynaptic marker Bassoon. The micrographs highlight the abnormal elongated shape of the Bassoon marker. The lower right insets represent color-coded Bassoon signal based on the size of the objects: below 0.8 μm^3^ (cyan) or above (purple). (**E-H**) Quantifications of the Bassoon signal from **D**. (**E**) The graph shows sphericity index of object above 0.8 μm^3^, highlighting that SARS-CoV-2 infection correlates to elongated presynaptic Bassoon structures. The n corresponds to the number of objects from one organoid. Two-tailed p value was < 0.0001 (****). (**F-G**) The graphs show the distribution of Bassoon object volumes for each condition. (**F**) Each dot corresponds to a single Bassoon object obtained from one organoid. Two-tailed p value was < 0.0001 (****). (**G**) The binning of the Bassoon objects by volume highlights a marked increase of the number of Bassoon objects of 2 μm^3^ and above. (**H**) Quantification of the percentage of Bassoon objects > 2 μm^3^. Each dot corresponds to fields of view acquired from three organoids. Two-tailed p value was < 0.01 (**). (**I**) Three-dimensional imaging of organotypic *ex vivo* culture of *post mortem* human frontal brain slices mock-infected or infected with SARS-CoV-2 at 10^5^ pfu/well for 4 days. Slices were stained for the presynaptic marker Bassoon. (**J**) Quantification as in G of the Bassoon signal from the culture of *ex vivo* brain slices shown in **I**. The binning of the Bassoon objects by volume highlights a marked increase of the number of Bassoon objects of 4 μm^3^ and above. (**K-P**) Analyses of the recording of the local field potential, recorded by the 3D-MEA, using Random Forest Classifier (RFC). (**K**) The discrimination accuracy between mock-infect and SARS-CoV-2-infected organoids at 0, 0.5 and 24 hpi. Here, 0.5 represents the chance level and 1.0 represents perfect discriminability. (**L**) The RFC highlighted two frequency bands-a lower and higher frequency band (highlighted by the black squares) in which the RFC found differences between the infected and mock organoids. The RFC looks at inter-frequency relations, and can thus find differences even when such are not visible on direct visual comparison, like in the lower band (**M**, **O**). However, in the higher band (**N**) we could see visibly different power spectrums for the infected and mock organoids (**P**).

In a second step, we reanalyzed our single organoid proteomic dataset using the Synapse Gene Ontologies software (SynGO) (*27*). Focusing on the 180 proteins upregulated upon infection with p values < 0.05 (no fold-change threshold applied), we found that about a third of them were part of the synaptosome (**Fig. 2B-C** and **Table S3**). Quantitative image analyses of the presynaptic marker bassoon in organoids highlighted that these structures were significantly enlarged and elongated upon exposure to SARS-CoV-2. (**Fig. 2D-H**). As organoids are surrogates for an immature brain with high neuronal plasticity, we next interrogated whether such phenotype could be observed in a model with low synaptic plasticity using our *ex vivo* organotypic brain slices (**Fig. 2I-J** and **Suppl Fig. S5A**). We found that bassoon object size was significantly upregulated upon infection, with similar elongated structures as seen in infected organoids (**Fig. 2D-H**). Moreover, exposure to UV-inactivated SARS-CoV-2 particles was sufficient to induce an increased number of trans-synaptic complexes in human primary neurons (**Suppl Fig. S5B-C**), further highlighting that virus replication is not required to trigger perturbations. Of note, enlarged synapses have been previously reported in the context of synaptic homeostasis upon chronic blockade of AMPARs and NMDARs, coined as “synaptic upscaling” (*28*).

### SARS-CoV-2 exposure perturbs local field potential

Neural synapses are subcellular structures essential for electrical signal transmission. Thus, we wondered whether SARS-CoV-2 could perturb electrical activity *in vitro*, as this has been reported in EEG from COVID-19 patients (*8, 9, 29*). We performed and compared local field potential recordings on cortical organoids in the presence or absence of SARS-CoV-2 using 3D MEAs (**Suppl Fig. S6A-B**). We developed a machine learning framework to specifically and unbiasedly investigate these features. Using Random Forest Classifier (RFC), we found that organoids exposed to SARS-CoV-2 could be readily discriminated from their mock counterparts based on local field potential (**Fig. 2K**). Our algorithm was able to predict whether an organoid was exposed to SARS-CoV-2 with > 97% accuracy and we confirmed that this was not due to organoid-to-organoid intrinsic heterogeneity (**Suppl Fig. S6C**). The same infected organoids could not be discriminated from the mock ones prior infection nor 30 min post infection, further confirming the specificity of our procedure. Moreover, we identified two frequency bands, a *low band* ([0-330] Hz) and a *high band* ([3800-4200] Hz), that were critical to discriminate between mock and SARS-CoV-2 infected organoids (**Fig. 2L-N** and **Suppl Fig. S6D-F**). While the differences in the spectral power in the low band region are not visible to the naked eye (**Fig. 2M, 2O**), the differences in the high band showed a clear peak, indicative of a significant increase of the electrical signal amplitude in SARS-CoV-2-exposed organoids compared to the mock (**Fig. 2N, 2P**). Together, our data indicates that SARS-CoV-2 induces abnormal synaptic structures associated to perturbed electrical synaptic transmission.

### SARS-CoV-2 particles associate and dwell at trans-synaptic sites

To gain further insight into the mechanism underlying this phenomenon, we went back to our single-organoid proteomic dataset analyzed with SynGO and focused on synaptic proteins with high and consistent upregulated expression upon infection (**Fig. 3A** and **Table S3**). Among these, Latrophilin-3 (LPHN3 protein from the *adgrl3* gene) caught our attention, as it is a post-synaptic G-protein coupled receptor (GPCR) involved in synapse formation and maintenance (*30–32*). We found that LPHN3 was also upregulated in human brain explants by western blot analyses and immunostaining (**Fig. 3B-C**). Interestingly, the protein FLRT3, the main receptor of LPHN3 (*33*), was recently identified to be closely associated to the viral protein Spike in a BioID proximity labeling assay (PLA) (*34*). However, we were not able to highlight interactions between FLRT3 and Spike in primary neurons by co-immunoprecipitation, while ACE-2 successfully pulled-down (**Suppl Fig. 7A**). The transient, indirect, and low-affinity interaction between Spike and FLRT3 could explain this result. Therefore, we next used PLA imaging to highlight that SARS-CoV-2 particles were indeed associated to FLRT3 structures (**Fig. 3D**). We reasoned that electrical perturbation and LPHN3 upregulation could be attributed to viral particles lodging at these trans-synaptic regions. Using neurons exposed to replication-competent SARS-CoV-2 particles (*35*), we observed an accumulation of virus at FLRT3-LPHN3 trans-synaptic sites (**Fig. 3D-E**). Over time, we observed that viral particles increasingly localized at sites containing pre/post-synaptic markers at 16 hpi (**Fig. 3F-I and Suppl Fig. 7B-D**), suggesting that particles could be retained onto synaptic structures. Using our fluorescent virions, we performed confocal 3D live cell imaging on neurons labeled with SynaptoRed, a dye enriched in synaptic regions. We observed that SARS-CoV-2 particles dwelled steadily for prolonged periods of time when associated to the synaptic marker (**Fig. 3J-L**), further confirming that SARS-CoV-2 particles can be retained at synapses.

**Figure 3.**
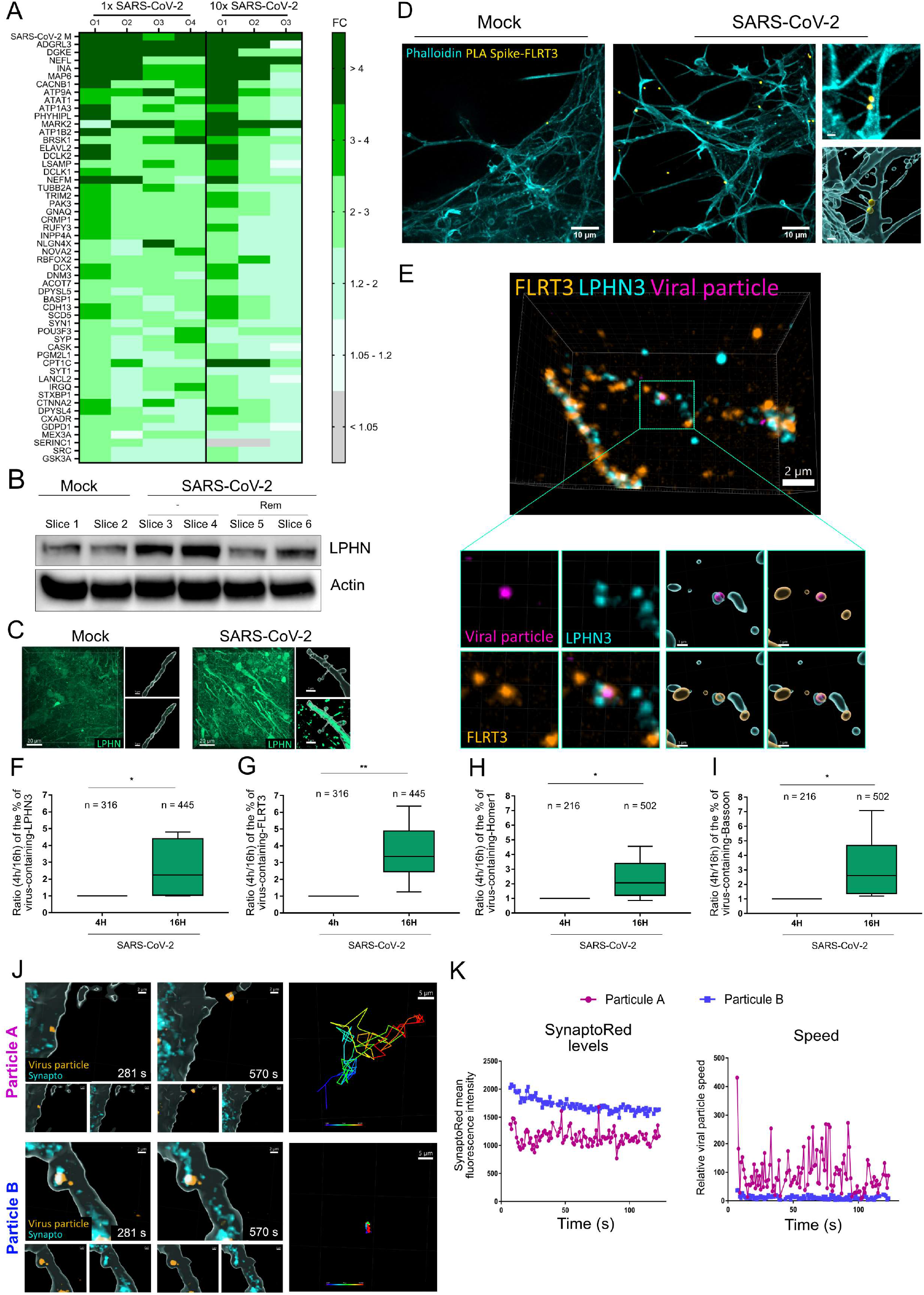
SARS-CoV-2 particles associate with Lphn3/FLRT3-containing synapses. (**A**) Heatmap of the upregulated proteins identified by single-organoid differential proteomics. The map shows the color-coded fold change (FC) of 1x and 10x SARS-CoV-2 exposure for each individual organoid. The most enriched protein in the infected samples (> 20 folds) is the viral Matrix (M), followed by the post-synaptic host protein ADGRL3 (Latrophilin-3/LPHN3), upregulated > 6 folds. (**B-C**) Organotypic *ex vivo* culture of *post mortem* human frontal brain slices mock-infected or infected with SARS-CoV-2 at 10^5^ pfu/well for 4 days. (**B**) The slices were individually lysed and immunoblotting analysis for LPHN3 and Actin as a loading control was performed. (**C**) Three-dimensional imaging of slices labeled with an anti-LPHN3 antibody. The insets highlight the shape of elongated cellular structures (upper panels) and the intensity of the LPHN3 signal (lower panels). (**D**) Proximity ligation assay (PLA) using anti-Spike and anti-FLRT3 antibodies and counterstained with an actin marker (Phalloidin). The yellow dots correspond to regions where FLRT3 and the viral protein Spike are closely associated. (**E-I**) Human neurons differentiated from neural progenitor cells (NPCs) were exposed to replication-competent SARS-CoV-2 particles harboring a mix of wild-type and fluorescent mRuby3-modified N protein for 4 h or 16 h. Cells were stained with LPHN3, FLRT3, Homer-1 and/or Bassoon as indicated. (**E**) Three-dimensional imaging and zoom-ins of SARS-CoV-2-exposed neurons shows a viral particle colocalizing with LPHN3 and FLRT3. (**F-I**) The graphs represent the percentage of virus particles containing the indicated synaptic marker at 16 h compared to 4 h post-infection. Data are mean +/- SD from at least six fields of view, with the n corresponding to the number of considered objects per condition. Two-tailed p value was < 0.05 (*) or < 0.01 (**). (**J-K**) Threedimensional live cell imaging of human neurons labeled with the synaptic marker SynaptoRed (Cyan) and exposed to SARS-CoV-2 N-mNeonGreen fluorescent particles (orange) for > 2 h. (**J**) The snapshots were extracted from two time-points from Movies S1 and S2, highlighting two viral particles and their associated single-particle tracking. (**K**) The graphs show SynaptoRed levels localized at the particles and viral particle speed from Movies S1 and S2. While Particle A (purple) has low levels of SynaptoRed and moves rapidly, Particle B (blue) is associated with high level of the synaptic marker and dwell on there for the all movie duration.

NeuroCOVID remains poorly understood at the molecular and subcellular levels. Although hypoxia and neuroinflammation have been described in patients with severe COVID-19, it cannot fully explain the persistence of subtle neurological and/or psychiatric disorders in the acute and post-acute phase of the disease. Because the CNS expresses low to undetectable levels of ACE-2, most neural cells are poorly supporting SARS-CoV-2 replication and dissemination, fostering investigations on alternative scenarios. Here, we discovered that SARS-CoV-2 dramatically modifies the synaptosome landscape and perturbs local field potential. These observations were associated to the retention of viral particles at the neural synapse, suggesting a mechanism of direct host-pathogen interference. Synaptic plasticity and maintenance are fine-tuned processes that are tightly regulated to quickly propagate information through electrical activity. A single grain of sand – a virion – in the gears, and the whole system gets jammed. By showing that the physical hindrance of viral particles at the synapse lead to profound perturbations and dysfunctions, we introduce a novel paradigm that is concordant with the observations made *in vitro, in vivo* and in patients.

## Acknowledgments

Image acquisitions in BSL-2 environment were performed at the MRI imaging facility (CNRS, Univ Montpellier), which also provided advice and training. All infections and live cell imaging using replication-competent SARS-CoV-2 were performed at the CEMIPAI BSL-3 facility (CNRS, Univ Montpellier).

## Funding

CNRS INSB (RG)

Agence Nationale de la Recherche ANR-20-CE15-0019-01 (RG)

Agence Nationale de la Recherche ANR-21-CE33-0007-03 (RG and GG)

CBS2 Montpellier doctoral school (EP)

FRM-ANRS MIE (RG and EP)

Agence Nationale de la Recherche ANR-10-INBS-08-03; ProFI FR2048 (CC)

Isite MUSE (Montpellier University of Excellence) (BC)

European Union’s Horizon 2020 Programme (H2020-INFRAIA-2018-1) [823839] (LM)

Research Foundation Flanders (FWO) [G028821N] (LM)

Research Foundation Flanders (FWO) [1S57123N] (TC)

Ghent University Concerted Research Action [BOF21/GOA/033] (LM).

Fondation pour la Recherche Médicale (FRM), MIE202207016212 (EP, RG)

## Author contributions

Conceptualization: RG

Methodology: EP, AH, WL, TC, FD, MSD, JB, LM, BC, CC, SC, GG, RG

Investigation: EP, AH, FD, WL, MSD, JB, DC, LA, ML, CC, SC, GG, RG

Analysis: EP, AH, FD, WL, TC, LM, MSD, JB, DC, LA, ML, CC, SC, GG, RG

Supervision: LM, CC, SC, GG, RG

Writing – original draft: RG

Writing – review & editing: EP, AH, TC, LM, MSD, DC, CC, SC, GG, RG

## Data and materials availability

All data are available in the main text or the supplementary materials.

## Supplementary Materials

### Materials and Methods

#### Antibodies and dyes

The following antibodies were used for immunofluorescence in this study: rabbit polyclonal anti-MAP2 (GeneTex), rat anti-CTIP2 (BioLegend), rabbit anti-Pax6 (BioLegend), goat anti-GFAP (Novus Biological), mouse anti-Tuj1 (Genetex), rabbit anti-Iba1 (Genetex), rabbit anti-N SARS-CoV-2 (Sino Biological), mouse anti-Spike SARS-CoV-2 (Genetex), mouse anti-Bassoon (Abcam), Rabbit anti-Homer1 (Synaptic system), Rabbit anti-LPHN3 (Proteintech), Rabbit anti-LPHN3 (ThermoFisher), Mouse anti-Lphn3 (Santa Cruz Biolotechnology), Goat anti-FLRT3 (R&D Systems), Rabbit anti-Cleaved Caspase 3 (Cell Signalling Technologies),

The following antibodies were used for histology examination: NeuroF (NF70, 1/100 clone 2F11, DAKO), Mouse anti-NeuN (1/400, clone A60, MAB377, Millipore), rabbit anti-Oligo2 (1/100, clone EP112, BSB2562, BioSB/diagomics), Mouse anti-GFAP (1/400, clone 6F2, MO761, DAKO).

The following antibodies were used for immunoblotting in this study: rabbit monoclonal anti-ACE-2 (Novus Biological), rabbit anti-Actin (Genetex), rabbit anti-N SARS-CoV-2 (Sino Biological), mouse anti-LPHN3 (Santa Cruz Biolotechnology).

The following dyes were used in this study: Dapi Nuclear Counterstain (Pierce), Hematoxylin and eosin (H&E), Live/Dead Viability/Cytotoxicity (ThermoFisher), SynaptoRed C2 (Biotium),

#### Stem cells culture and cortical organoid differentiation

The human embryonic stem cells (hESCs) H9 cultured and differentiated into cortical organoids as in (*36*). Briefly, hESCs were cultured on 60 mm Matrigel (corning) coated plate in mTeSR PLUS media (StemCell Technologies). Cells were split by squaring colonies with a needle and 3 colonies were used for a 60 mm plate. For organoid production, colonies were split onto a 100 mm plate with mitomycin-C-treated MEF feeding layer cells in DMEM/F12 supplemented with Glutamax (Gibco) with 20% KnockOut serum replacement (Gibco), 1x non-essential amino acids (Gibco), 100 μM β-mercaptoethanol (Gibco), and 1x Penicillin–Streptomycin (Gibco). The media is changed every day with supplementation with 8 ng/ml human βFGF (Sigma-Aldrich). Colonies were scooped with a cell lifter (Corning) and cultured in low binding 60 mm dish in differentiation media (hESC media supplemented with 1x sodium pyruvate). The next day, hESC differentiation was initiated by replacing half of the media with media containing 2x of the following small molecules: 3μM IWR-1-Endo, 1 μM Dosomorphin, 10 μM SB-431542 and 1 μM Cyclopamine, all obtained from Sigma-Alrdich. Media was renewed every other day. At day 18 postdifferentiation, the growing organoids were cultured in N2 media, composed of Neurobasal media (Gibco), 1x N2 supplement (Gibco), 2 mM L-glutamine (Gibco), 1x Penicillin–Streptomycin and 1 μM Cyclopamine. At day 24, the Cyclopamine was remove from the media and organoids were grown at least until day 35 to 45 post-differentiation before used for experiments.

#### Post mortem brain explant and ex vivo organotypic culture

The protocol for post mortem brain explant was accepted by the institutional review board (IRB) of the CHU of Montpellier and French Biomedicine Agency, permitting human brain resection for research purpose during autopsy performed at the forensic institute of the CHU de Montpellier, and family consent was obtained. All the data presented originate from 4 patients aged from 25 and 72 years old. Patients were chosen based on the absence of cerebral injury or trauma and the post-mortem interval (PMI) was not exceeding 12 h. The frontal and parietal regions from the motor cortex were first dissected by making several 1-3 cm-thick coronal incisions of the brain, from frontal to occipital region and second, by cutting small cubes (about 0.5 cm^3^) perpendicularly to the longitudinal axis of the gyrus and kept in N2 media at 4°C before further processing. The cubes were embedded in 3% low melting point agarose (ThermoFisher) and sliced in 300 to 600 μm thick sections using an PELCO easislicer (Ted Pella). All the samples display cortical layers and a piece of the white matter. The brain slices were dissociated from agarose and transferred onto cell culture inserts with 0.4 μm pore PET membranes (Millicell, Millipore) hanging in 6-well plates containing 2.5 ml of N2 media. The slices were cultured at 37°C, 5% CO_2_ and 95% humidity at the air-liquide interface (ALI) and media was changed twice a week (see also **Suppl Fig. S2**).

#### COVID19 patients

The research protocol (Academic Hospital and University of Reims) was approved by the French Biomedicine Agency (n°PFS20-022). Family consent was obtained, and absence of opposition from the patient was checked at the French Biomedicine Agency for each patient. All patients (six) had a positive nasal swab for SARS-CoV-2 before death, and were hospitalized in Intensive Care Unit with a severe COVID19 diagnosis, between May of 2020 and May of 2021. Post mortem delay before samples was less than 24 hours. Four male and two female patients were sampled, with a median age of 64.5-year-old (44 to 79-year-old). Delay between COVID19 diagnosis and death was 11 to 32 days. Blood (sus-clavicular or femoral veins), and cerebro-spinal fluid (lumbar puncture) were taken when possible. Samples of temporal regions of the brain lobes were immediately frozen at −80°C, and then the brain was fixed in formaldehyde 4%. Frontal and parietal regions were sampled and embedded in paraffin. RNA samples were extract using the KingFisher Flex systems (ThermoFischer) according to manufacturer’s instructions. RT-qPCR was performed on 5 μl of RNA using the Quantabio ToughMix reagents (Quantabio). The IP2/IP4 probe sets (*37*) were used to detect SARS-CoV-2 RNA. GAPDH mRNA was also checked. PF-CRB of the Academic Hospital of Reims stored and provided the samples for analyses at the CHU of Montpellier (n°2022-MAD30).

#### Neural stem cell line culture and neuron differentiation

Human neural progenitor cells (hNPCs), were obtained from Creative Biolabs (NCL-2103-P83). These cells originate from induced pluripotent stem cells (iPSCs), obtained from adult skin fibroblasts. NPCs were cultured in flasks coated with Matrigel (Corning) with STEMdiff Neural Progenitor Medium (StemCell Technologies). Differentiation was performed on coverslips coated with 100 μg/ml poly-L-ornithine and 10 μg/ml laminin (Merck) in differentiation media (50% DMEM/F-12 + Glutamax, 50% Neurobasal, 1x N2, 1x B27). The media was changed every other day. After eight days, 10 μg/mL BDNF and 10 μg/mL GDNF were added to the media every day until day 14, on which differentiated neurons were used for experiments.

#### Virus production

The virus used in this study is the BetaCoV/France/IDF0372/2020 isolate, and was supplied by Pr. Sylvie van der Werf and the National Reference Centre for Respiratory Viruses hosted by Institut Pasteur (Paris, France). The patient sample from which strain BetaCoV/France/IDF0372/2020 was isolated was provided by Dr. X. Lescure and Pr. Y. Yazdanpanah from the Bichat Hospital, Paris, France. The BetaCoV/France/IDF0372/2020 was amplified in Vero E6 cells at MOI 0.001 and supernatant was harvested at 72 hpi (when about 50% cytopathic effects were observed). Cell debris were removed by centrifugation at 2000 x g for 5 min, and aliquots stored at −80°C. The titer was defined by plaque assays in Vero E6 cells. Typical titers were ranging around 10^7^ pfu/ml. SARS-CoV-2 mNeonGreen and NLuc reporter viruses, in which the ORF7 of the viral genome was replaced with the indicated reporter gene, were obtained from (*22*). These replication-competent viruses express the reporter gene upon entry, replication and translation of the virus in receiving cells. The SARS-CoV-2 N-mNeonGreen particles were generated as in (*35*). Briefly, Vero E6 cells, stably expressing N-mNeonGreen from SARS-CoV-2 (Addgene # 170467) were infected with wild-type SARS-CoV-2 at MOI 0.001. Cells were washed 24 hpi in PBS, the supernatant was harvested at 72 hpi and debris were removed by centrifugation at 2000 x g for 5 min. UV-inactivated virus was obtained by incubating 250 μl of viral preparation in a UV-C 254 nm UV chamber (Vilber Lourmat Biolink BLX UV Crosslinker) at 2 joules. The loss of infectivity was confirmed by infection of Vero E6 cells. Mock-infected samples correspond to supernatant of non-infected Vero E6 cells processed as infected cells. Remdesivir (MedChem Express) was used at 10 μM to inhibit virus replication. NLuc luminescence was measured using the Nano-Glo Luciferase Assay System (Promega) and read by PerkinElmer EnVision spectro-luminometer.

#### Immunohistochemistry

After formalin fixation and paraffin embedding, 5-μm sections were stained with hematoxylin and eosin. Immunohistochemistry was performed with Autostainer BenchMark ULTRA (Ventana Medical Systems), according to manufacturer’s instructions. Briefly, paraffin sections were incubated at 60°C for 16 min and then at 72°C for dewaxing and at 95°C in retrieval antigen buffer. Incubation for 30 min with antibodies was performed and revealed with DAB. Samples were washed and incubated with Hematoxylin II (counterstaining) for 12 min and with bluing reagent (counterstaining) for 4 min.

#### Immunostaining of organoids and brain slices

Samples were washed with phosphate buffered saline (PBS) once and transferred in 96-well plate round bottom (for brain organoids) or in 24-well plate flat bottom (for brain slices). Samples were fixed in 4% paraformaldehyde (PFA) for 1 h at room temperature (RT). Samples were washed once with PBS and permeabilized for 24 h on a rocker with permeabilization buffer (0.5% BSA, 1% Triton in PBS). All subsequent steps were performed in permeabilization buffer. Samples were labeled with primary antibody for 24 h on a rocker, washed overnight, labeled with appropriate secondary antibodies for 24 h and washed overnight. The samples were incubated RapiClear 1.52 reagent (Sunjin Lab) overnight at RT.

#### Fluorescence confocal microscopy

Image acquisition was performed on a spinning-disk confocal microscope (Dragonfly, Oxford Instruments) equipped with an ultrasensitive 1024 × 1024 EMCCD camera (iXon Life 888, Andor) and four laser lines (405, 488, 561, and 637 nm). A 20x, NA 0.8 air objective was used for whole sample imaging and a 60x, NA 1.15 (Nikon) oil-immersion long-distance objective was used for in-depth imaging of selected areas. Images were processed using FiJi (ImageJ) and Bitplane Imaris x64 (Oxford Instruments) version 9.2 and 9.7.

#### Live cell imaging

Neurons were differentiated on a 30-mm diameter #1.5 glass coverslip in six-well plates. Before imaging, the cells were incubated with SARS-CoV-2 N-mNeonGreen in neuronal media. Image acquisition was performed at 37°C and 5% CO_2_ in a dark chamber using an AxioObserver.Z1 inverted microscope (Zeiss) mounted with a spinning disc head (Yokogawa), a back-illuminated EMCCD camera (Evolve; Photometrics), and a 100×, 1.45 NA oil objective (Zeiss) controlled by MetaMorph (Molecular Devices) software.

#### Image analysis and quantification

Processing and image quantification were done using Bitplane Imaris v9.7. Bassoon object were identified using absolute intensity thresholding and converted to a surface rendering model. The parameters used in this study (sphericity, volume, intensity) were automatically extracted from each object identified by intensity thresholding upon background subtraction. For virus-synapse association studies, each object was identified upon absolute intensity thresholding and converted using the “Spot” feature on the Imaris software. a region comprised between 0 and 0.5 μm from the spot center was used to consider that a virus particle was associated with synaptic proteins. Data were extract automatically to excel files format. Graphs and statistics were performed using GraphPad Prism version 9.

#### Western blot

Samples were lysed in RIPA buffer (150 mM sodium chloride, 1% NP-40, 0.5% sodium deoxycholate, 0.1% SDS, 50 mM Tris pH 8.0) supplemented with protease inhibitor (Promega). A pellet pestle (VWR) was used to favour protein extraction from tissues. Lysates were kept at −80°C prior analysis. Samples were spun down at 15 000 x *g* for 15 min at 4°C, the supernatant was collected and protein concentration was measured using a Pierce BCA Assay Kit (Pierce). A total of 10 to 20 μg of protein lysate was run on Bolt 4 to 12% Bis-Tris plus gels (ThermoFisher), and proteins were transferred to nitrocellulose membranes using iBlot2 (ThermoFisher). Nitrocellulose membranes were blocked with 5% (w/v) milk in PBST (PBS pH 7.4, 0.05% Tween 20) for 30 min. Primary antibodies were incubated for 1 h at RT or overnight at 4°C in PBST containing 5% milk. Secondary antibodies were incubated for 1 h at RT. After washes with PBST, nitrocellulose membranes were incubated with Clarity Max Western ECL Substrate (Bio-Rad). The specific proteins were visualized with a ChemiDoc imaging system (Bio-Rad).

#### Immunoprecipitation

Protein lysates from neurons were incubated overnight at 4°C with 5 μg of mouse anti-Spike antibody (Genetex). The next day, 20 μl of protein G agarose beads (Pierce) were added to the lysate and incubated at 4°C for 2 h. The beads were spun down at 3000 x g for 3 min and washed three times with PBS. The sample was eluted in NuPAGE LDS sample buffer (Thermofisher) in presence of 50 mM DTT.

#### RT-qPCR

RNA samples were extract using the RNeasy KIT (Qiagen) according to manufacturer’s instructions. RT-qPCR was performed on 2 μl of RNA using the TaqPath One-Step RT-qPCR, CG master mix (ThermoFisher). The N1 primer/probe sets designed by the Center for Disease Control (CDC) were used to detect SARS-CoV-2 RNA. All samples were normalized to RPL27 mRNA.

#### MS-based quantitative proteomics

##### Sample preparation

Individual brain organoids were lysed in Laemmli buffer composed of 10 mM Tris pH 6.8, 1 mM EDTA, 5 % SDS and 10 % glycerol. Samples were sonicated for 2 min. Protein concentration of all samples was determined using the DC protein assay (Bio-Rad) according to manufacturer’s instructions, in triplicate and a standard curve was established using BSA. Dithiothreitol (DTT) was added at a final concentration of 50 mM and cell lysates were heated at 95°C for 5 min. Protein extracts were stacked in an in-house prepared 5% acrylamide SDS-PAGE stacking gel at 50 V. Proteins in the gel were fixed with 50% ethanol/3% phosphoric acid, washed and colored with Silver Blue. Gel bands were cut, washed with ammonium hydrogen carbonate and acetonitrile, reduced with 10 mM DTT and alkylated using 55 mM iodoacetamide prior to overnight digestion at 37°C with modified porcine trypsin (Promega) at a final trypsin/protein ratio of 1/50. Trypsin-digested peptides were extracted with 60% acetonitrile in 0.1% formic acid followed by a second extraction with 100% acetonitrile. Acetonitrile (ACN) was evaporated under vacuum and the peptides were resuspended in H20 and 0.1% formic acid (FA) before nanoLC-MS/MS analysis.

##### NanoLC-MS/MS analysis

NanoLC-MS/MS analyses were performed on a nanoAcquity UltraPerformance Liquid Chromatography device (Waters Corporation) coupled to a quadrupole-Orbitrap mass spectrometer (Q-Exactive HF-X, Thermo Fisher Scientific). Peptide separation was performed on an ACQUITY UPLC Peptide BEH C18 Column (250 mm x 75 μm with 1.7 μm diameter particles) and an ACQUITY UPLC M-Class Symmetry C18 Trap Column (20 mm x 180 μm with 5 μm diameter particles; Waters Corporation). The solvent system consisted of 0.1% FA in water (solvent A) and 0.1% FA in ACN (solvent B). Samples (800 ng) were loaded into the enrichment column over 3 min at 5 μl/min with 99% of solvent A and 1% of solvent B. Chromatographic separation was conducted with the following gradient of solvent B: from 2 to 25% over 53 min, from 25 to 40% over 10 min, from 40 to 90% over 2 min.

The mass spectrometer was operated in data-dependent acquisition mode. Survey full scan MS spectra (mass range 300-1800) were acquired in the Orbitrap at a resolution of 60K at 200 m/z with an automatic gain control (AGC) fixed at 3 10^6^ and a maximal injection time set to 50 ms. The ten most intense peptide ions in each survey scan with a charge state ≥ 2 were selected for fragmentation. MS/MS spectra were acquired at a resolution of 15K at 200 m/z, AGC was set to 1 10^5^, and the maximal injection time was set to 50 ms. Peptides were fragmented by higher-energy collisional dissociation with a normalized collision energy set to 27. Peaks selected for fragmentation were automatically included in a dynamic exclusion list for 60 s. All samples were injected using a randomized and blocked injection sequence (one replicate of each group in each block). To minimize carry-over, two solvent blank injections were performed after each sample.

##### Data interpretation

Raw files were converted to .mgf peaklists using MsConvert and were submitted to Mascot database searches (version 2.5.1, MatrixScience, London, UK) against a human (20 342 sequences, 2021-04-20, Taxonomy ID 9606) and SARS-CoV-2 (17 sequences, 2021-04-20, Taxonomy ID 2697049) protein sequences database downloaded from UniProtKB-SwissProt, to which common contaminants and decoy sequences were added. Spectra were searched with a mass tolerance of 10 ppm in MS mode and 0.07 Da in MS/MS mode. One trypsin missed cleavage was tolerated. Carbamidomethylation of cysteine residues was set as a fixed modification. Oxidation of methionine residues and acetylation of proteins n-termini were set as variable modifications. Identification results were imported into the Proline software (version 2.6.2) (*38*) for validation and label-free quantification. Peptide Spectrum Matches (PSM) with pretty rank equal to one were retained. False Discovery Rate (FDR) was then optimized to be below 1% at PSM level using Mascot Adjusted E-value and below 1% at Protein Level using Mascot Mudpit score. For label free quantification, extracted ion chromatograms were used to derive peptides abundances. An m/z tolerance of 10 ppm was used. Alignment of the LC-MS/MS runs was performed using Loess smoothing. Cross assignment of peptide ions abundances was performed among the samples and controls using a m/z tolerance of 10 ppm and a retention time tolerance of 42 s. Protein abundances were computed using the sum of the unique peptide abundances normalized at the peptide level using the median.

To be considered, proteins must be identified in all three or four replicates in at least one condition. The imputation of the missing values and differential data analysis were performed using the opensource ProStaR software (*39*). Imputation of missing values was done using the approximation of the lower limit of quantification by the 2.5% lower quantile of each replicate intensity distribution (“det quantile”). A Limma moderated t-test was applied on the dataset to perform differential analysis. The adaptive Benjamini-Hochberg procedure was applied to adjust the p-values and False Discovery Rate.

The mass spectrometry proteomics data have been deposited to the ProteomeXchange Consortium via the PRIDE partner repository (*40*) with the dataset identifier PXD036485.

#### Post-translational modification analysis

Raw mass spectrometry files were converted to mgf format using ThermoRawFileParser v1.3.4 (*41*) and searched against the human complement of the UniProtKB/SwissProt protein database containing 20,397 proteins, with added common contaminants and 17 proteins from the Severe acute respiratory syndrome coronavirus 2 proteome (UP000464024; June 2022). The search was performed using the ionbot search engine (v0.7.0) (*42*) with open modification search settings and three expected modifications: (i) carbamidomethylation of cysteine; (ii) oxidation of methionine; (iii) and N-terminal acetylation. Results were filtered on 1% FDR using a q-value below 0.01. A global comparison of modification levels was calculated as the ratio of ratios. In infected samples, for each modification the ratio of the number of spectra of modified versus unmodified peptides was calculated and then divided by the same ratio in the mock samples. Relative quantification was done in two steps. First, a filtering step removed PTMs that accounted for less than 3% of all PSMs detected in the sample. Second, the filtered PTMs were quantified by dividing the number of PSMs matching this PTM, by all possible modification sites of the PTM in the sample. This fraction was then normalized by the sum of all these fractions in the sample.

#### Electrical activity recording

Cortical organoids were plated on 3D MEA (MultiChannel System), made of and array of 60 TiN-coated conical electrodes of 12 μm diameter active site of 80 μm height. The specific sharp shape of these electrodes allows a better penetration of the tissue than a planar MEA and the measurement of extracellular local field close to active cells. The surface of the MEA was pre-coated with 100 μg/ml poly-L-ornithine and 10 μg/ml laminin (Merck). After 2 days, electrical recording of the local field potential was performed inside an incubator at 37°C, 5% CO_2_ and 95% humidity using the MEA2100-Mini headstage (MultiChannel System) under electromagnetic shielding. Traces of electrical signals versus time were recorded at a sampling rate of 50 kHz by the Multichannel experimenter software with a customized bandpass filter script for 1 min per condition.

#### Electrical activity analysis using machine learning

Electrical activity was collected using a 60 channel MEA. The collected raw data from the electrodes were high pass filtered over 50 Hz and then sampled at 10000 Hz. We divided electrical recording on each day from the infected and non-infected (mock) organoids into 30-time bins each and performed frequency analysis on each bin. The frequency spectrum was sampled into 300 frequency features. We used the features from 70% of the recordings to train a Random Forest Classifier to detect an infected recording time-bin from a mock organoid time bin, and used it to test the detectability on the remaining 30% of the recordings. This process was repeated 10 times for each organoid on each day to determine the discriminability with each organoid, for that day. The top 5% of features isolated by RFC were observed to be bunched in two frequency bands, that were defined as the ‘low band’ and ‘high band’ respectively. The average of the frequency power within this bands were calculated for each organoid.

## Supplemental Figures S1 to S7

**Supplemental Figure S1.**
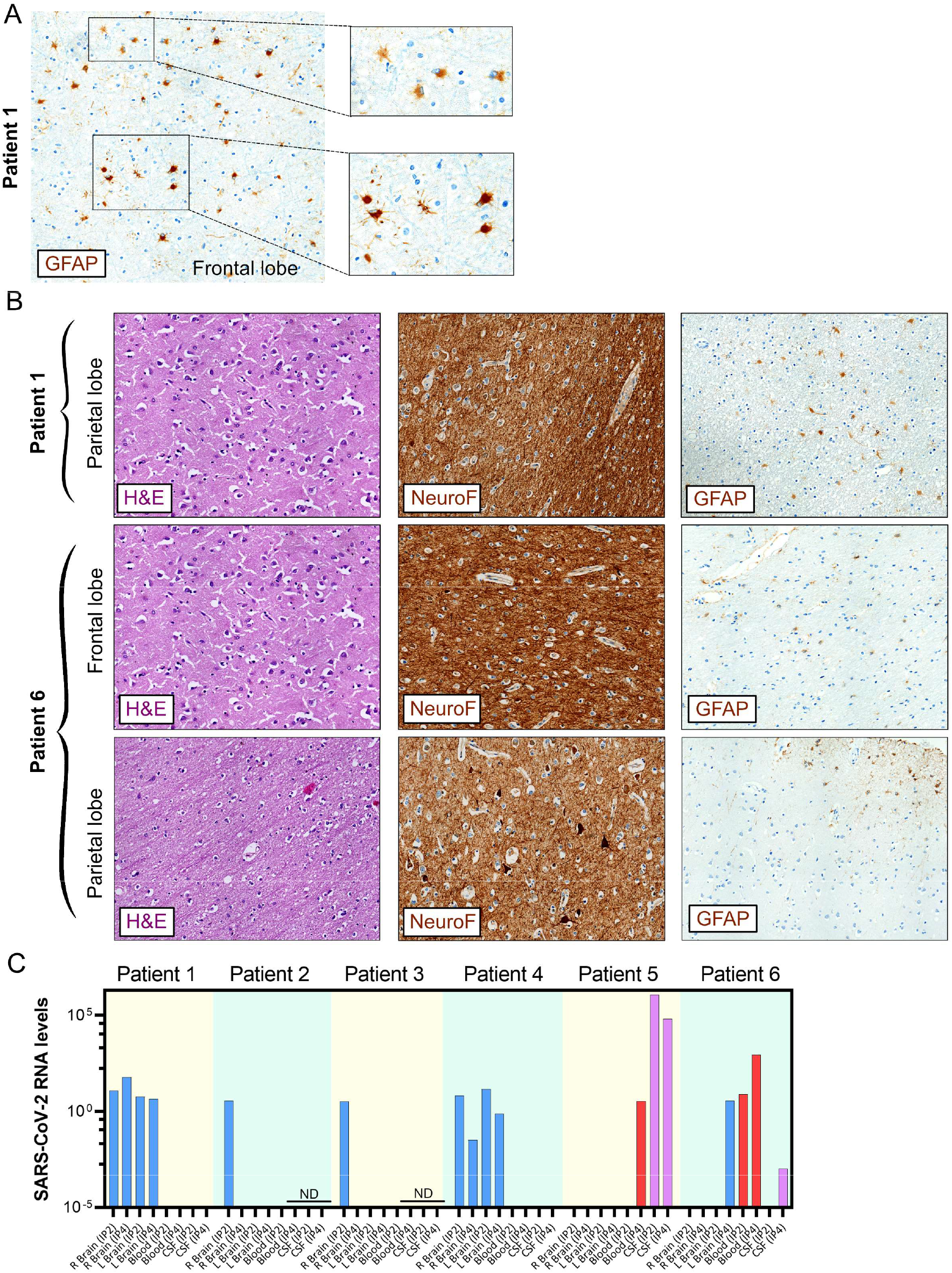
Characterization of the brain features of COVID-19 patients without neurological symptoms. (**A-B**) Imaging of Hematoxylin and Eosin (H&E), NeuroF and GFAP stainings using 20X magnification. (**C**) RT-qPCR of tissues obtained from the temporal lobe of the right (R Brain) and left (L Brain) hemispheres, blood and cerebrospinal fluid (CSF) using two anti SARS-CoV-2 probe sets (IP2 and IP4). GAPDH was also acquired. Color coding shows brain samples in blue, blood in red and CSF in purple. ND: not determined.

**Supplemental Figure S2.**
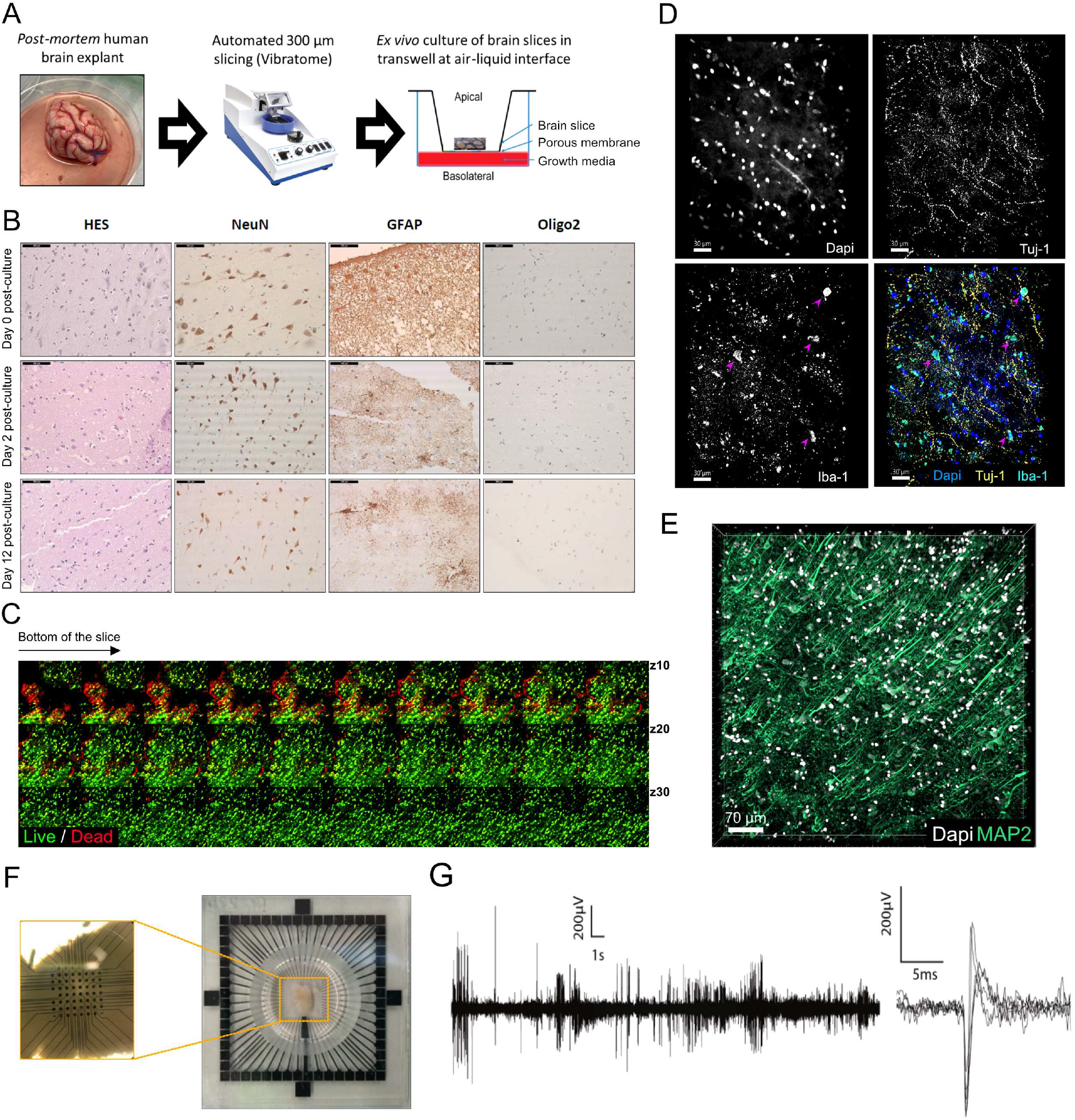
*Ex vivo* organotypic culture of post-mortem human brain explants. (**A**) Experimental procedure. (**B**) Imaging of Hematoxylin and Eosin (H&E), NeuN, GFAP and oligo2 stainings using 20X magnification. (**C**) Human brain slices from the frontal lobe were cultured for 17 days and stained with the Live/Dead marker, showing healthy cells (green), dead cells (red), and dying cells in yellow. The micrograph shows a gallery of confocal Z plans, spaced by 1 μm, from the bottom of the slice and getting deeper into the slice, showing healthy cells away from the cutting region. (**D-E**) Three-dimensional imaging of neurons (Tuj1 and MAP2) and microglial cells (Iba1) from a frontal slice cultured for 5 days *ex vivo* either non-infected (D) or exposed to SARS-CoV-2 (**E**). The neurons from SARS-CoV-2-exposed brain slices retain densely aligned organization. (**F-G**) Frontal brain slices were seeded onto 3D MEA (F) and electrical activity was recorded. (**G**) Example trace of extracellular electrical signal as a function of time recorded from one electrode and overlay cutouts showing the reproducibility of shape, amplitude, signal to noise ratio and dynamics of typical signals.

**Supplemental Figure S3.**
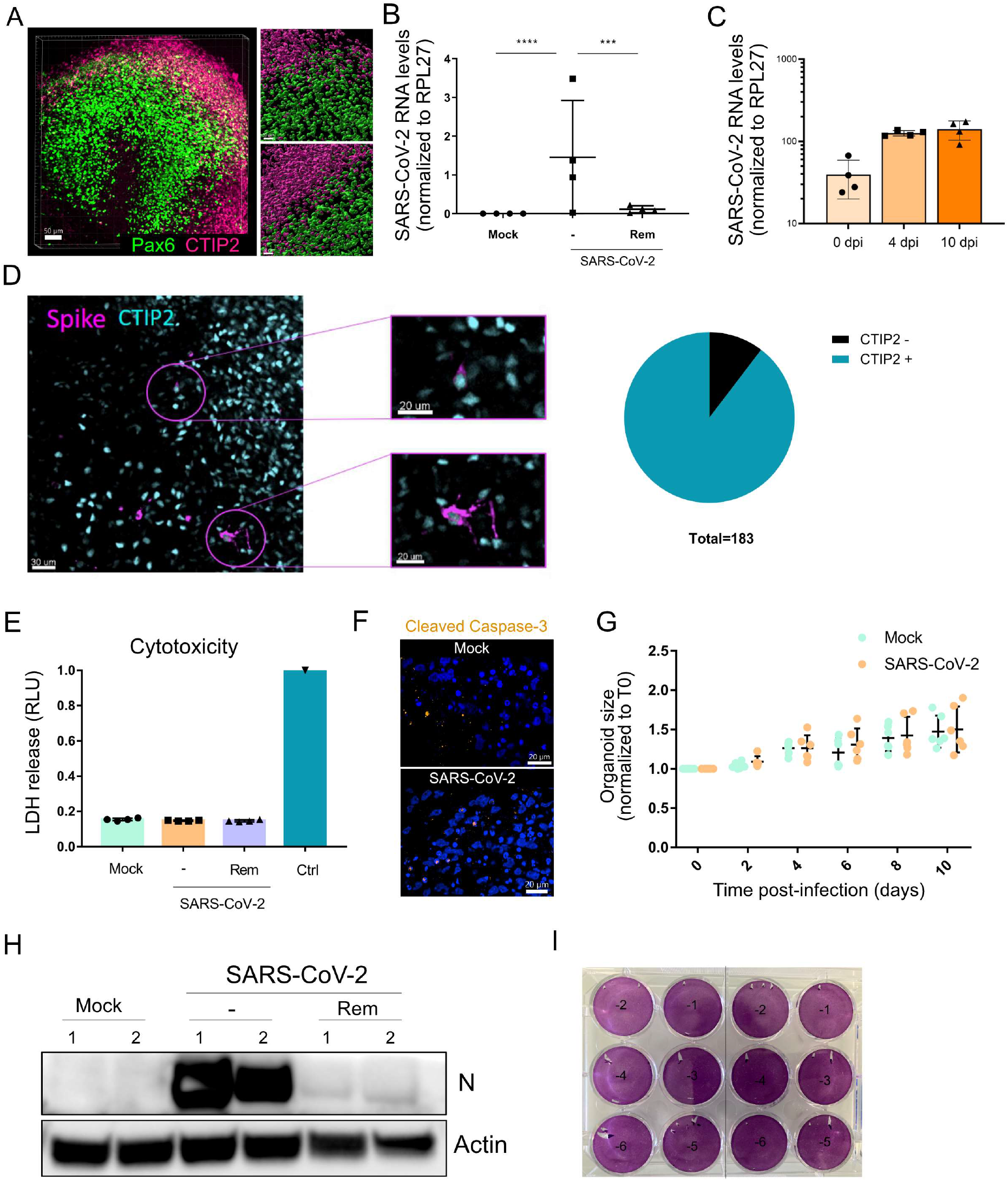
Characterization of SARS-CoV-2-exposed cortical organoids. (**A**) Three-dimensional imaging of a cortical organoid stained for neural progenitor (Pax6, green), and neurons (CTIP2, purple). (**B-C**) Cortical organoids were mock-infected or infected for 10 days. (**B**) or at indicated time (**C**) with SARS-CoV-2 at 10^5^ pfu/organoid in the presence or absence of 10 μM Remdesivir. RNA extraction and RT-qPCR were performed and the data corresponds to SARS-CoV-2 RNA levels normalized by RPL27 housekeeping gene from 2 organoids per condition in duplicates. Two-tailed p value was < 0.001 (***) or < 0.0001 (****). (**D**) Cortical organoids were infected with SARS-CoV-2 at 10^5^ pfu/organoid for 10 days, fixed and stain for Spike and the neuronal marker CTIP2. The micrograph and insets show CTIP2-positive infected cells. The pie chart made from 3 fields of view shows that ≈ 90% of the infected cells were positive for CTIP2. (**E**) Cortical organoids were mock-infected or infected with SARS-CoV-2 at 10^5^ pfu/organoid for 10 days in the presence or absence of 10 μM Remdesivir. LDH release was measured in the supernatant of the organoids. The data are from two organoids per condition in duplicates and shows no toxicity in any condition, except for Ctrl, a positive control used to account for maximum cytotoxicity. (**F**) Cortical organoids were mock-infected or infected with SARS-CoV-2 at 10^5^ pfu/organoid, fixed, permeabilized and stained with the anti-cleaved Caspase 3 antibody (orange, a cell death marker, and Dapi (blue). The micrographs show marginal apoptotic bodies in both conditions. (**G**) Cortical organoids were mock-infected or infected with SARS-CoV-2 at 10^5^ pfu/organoid, and their size was measured every 2 days for 10 days. Each dot corresponds to a single organoid. (**H**) Organoids processed as in **E**, were then lysed and western blot analysis was perform using anti-N and anti-Actin (loading control) antibodies. (**I**) the supernatant of samples processed as in **E**, was used for titration by plaque assay on Vero E6 cells. The labels −1 to −6 correspond to log10 dilution factors. The plate shows no plaques at any dilution.

**Supplemental Figure S4.**
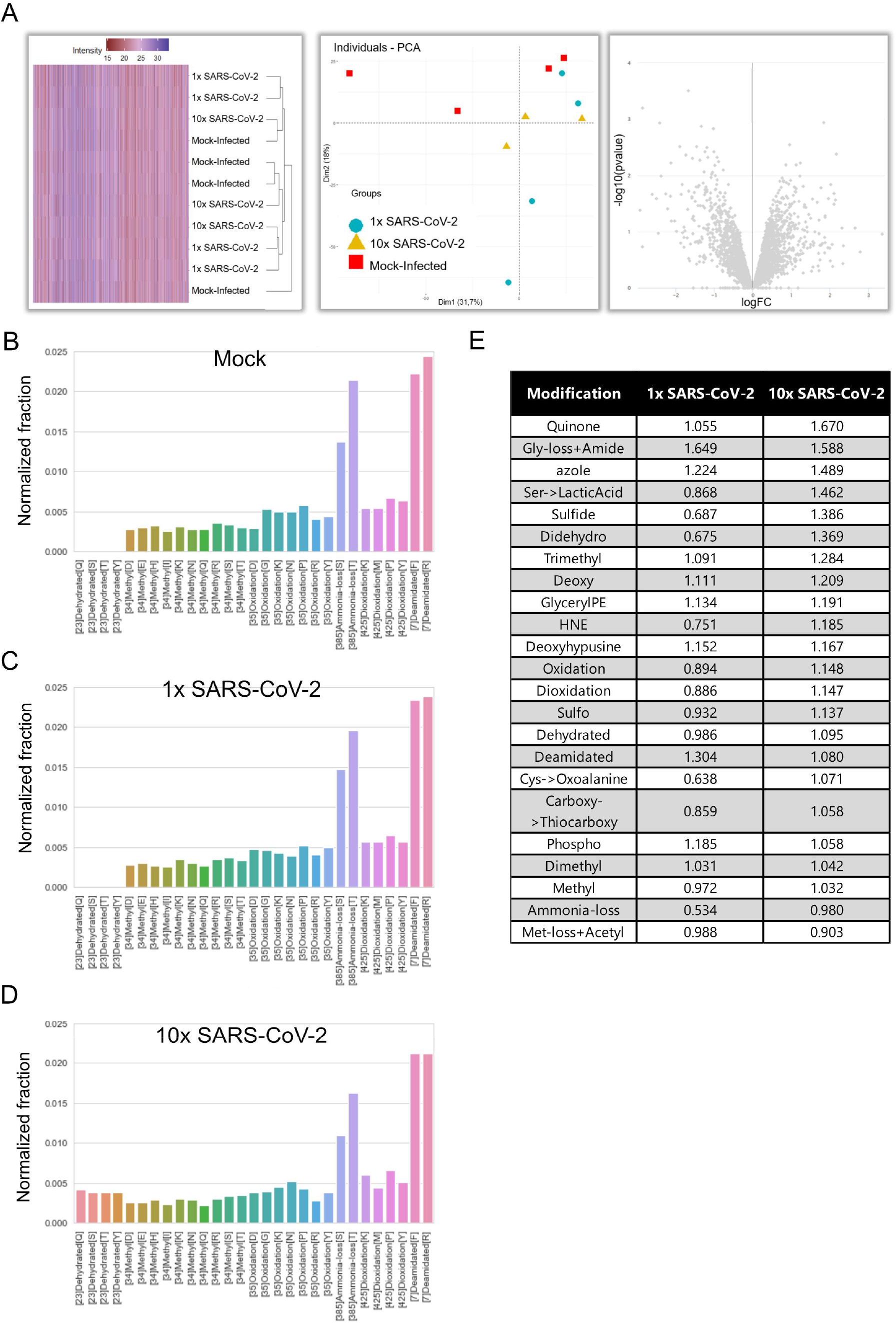
Single-organoid proteomic analysis. (**A**) Overview of the differential proteomic analysis results. Heatmap, clustering and principal component analysis on protein intensities obtained by summing unique peptide abundances and volcano plot representing foldchanges and p-values for all quantified proteins. (**B-F**) Patterns of unexpected post-translational modifications (PTM) in mock-infected, 1x infected and 10x infected samples obtained by averaging relative quantifications. **(E**) Global comparison of post-translational modification levels between infected and mock-infected samples by dividing the peptide modification ratio of the infected samples by the peptide modification ratio of the mock-infected samples. A value of 1 indicates no difference in modification ratio between samples, a higher value indicates an increased modification ratio in the infected samples and a value below 1 indicates a decreased modification ratio in the infected samples.

**Supplemental Figure S5.**
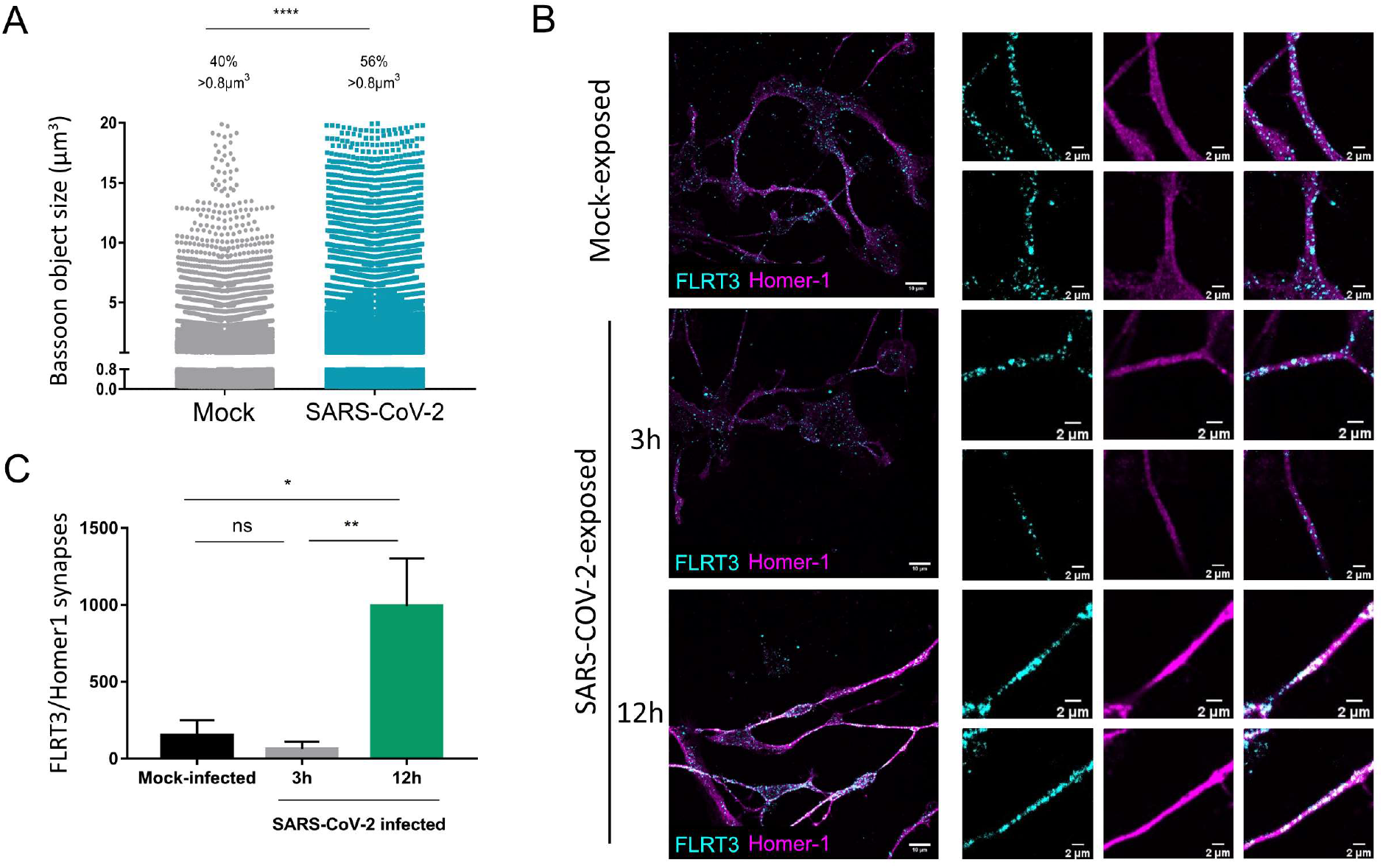
SARS-CoV-2 exposure impacts neural synapses. (**A**) The graph shows the distribution of Bassoon object volumes mock-infected or SARS-CoV-2-exposed human frontal brain slices at 10^5^ pfu/well for 4 days. Each dot corresponds to a single Bassoon object obtained from a two brain slices from two different donors. Two-tailed p value was < 0.0001 (****). (**B-C**) Mature neurons derived from hNPCs were mock-exposed or exposed to UV-inactivated SARS-CoV-2 for 3 or 12 h. Samples were fixed and stained for the presynaptic protein FLRT3 and the post-synaptic marker Homer-1. Image quantification shows a significant increase of the number of FLRT3/Homer-1 synapses at 12 h post-exposure. Two-tailed p value was < 0.05 (*) and < 0.01 (**).

**Supplemental Figure S6.**
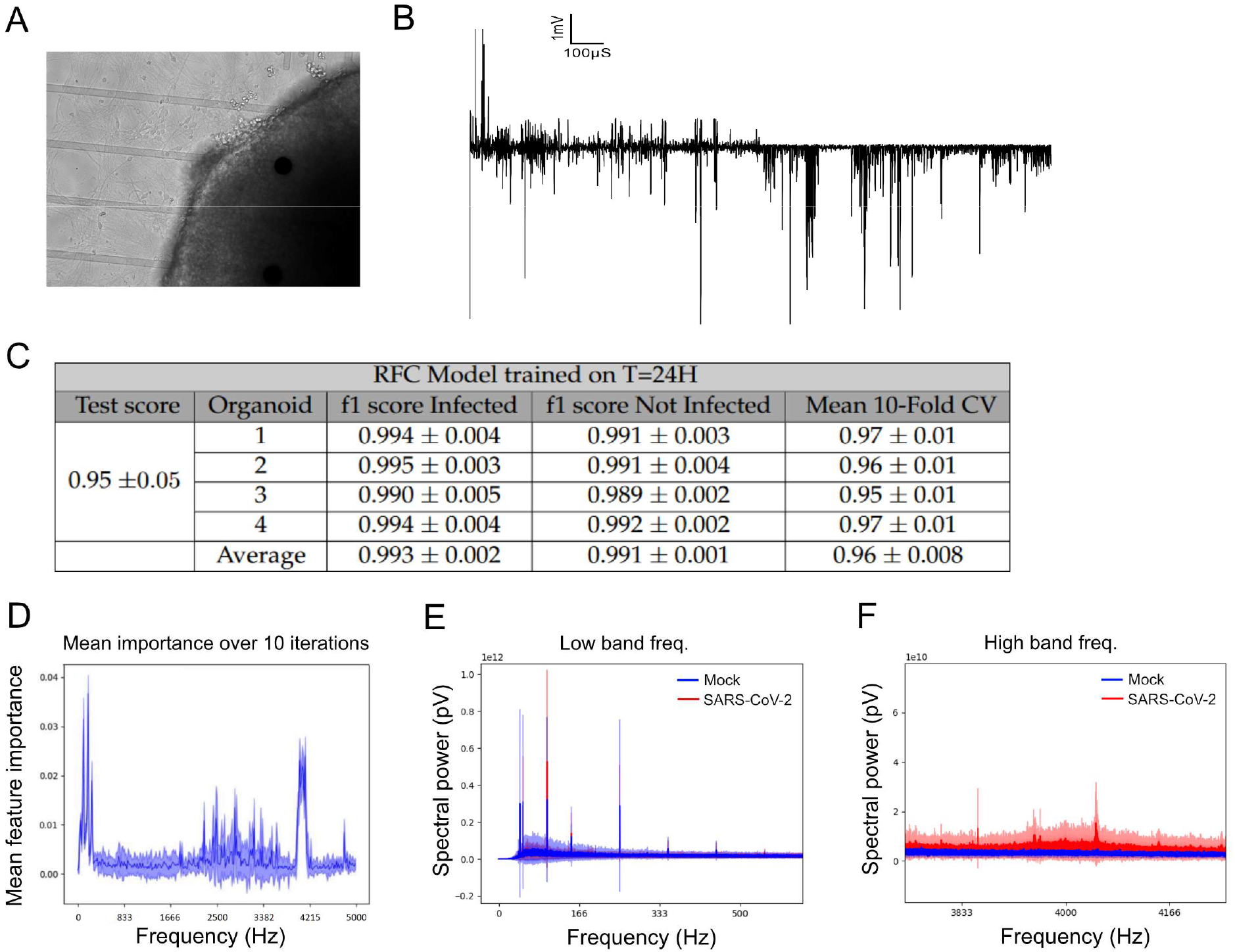
Exposure to SARS-CoV-2 impacts electrical activity. (**A**) Micrograph of an organoid on a 3D MEA electrode. (**B**) Extracellular electrical trace of an electrode on which a SARS-CoV-2-exposed cortical organoid was seeded. (**C)** A Random Forest Classifier (RFC) was trained on part of the infected and mock samples from all the organoids and tested on the untrained samples from each organoid. The table shows the probability of the RFC to classify an infected sample correctly (f1-score Infected) from each organoid, and the probability of the RFC to classify a non-infected sample correctly (f1 -score Non-Infected) from each organoid. **(D)** The plot of the frequency (feature) importance in the RFC classification. A higher value shows larger importance of the frequency in the discrimination of the infected and mock organoid. Note that lower two high importance frequency ranges (the low and high range). **(E**, **F)** Raw frequency spectrum plots of the electrical activity from the 4 organoids in low and high frequency range respectively. The smoothed version of the same plots are presented in Fig. 2M and 2N respectively.

**Supplemental Figure S7.**
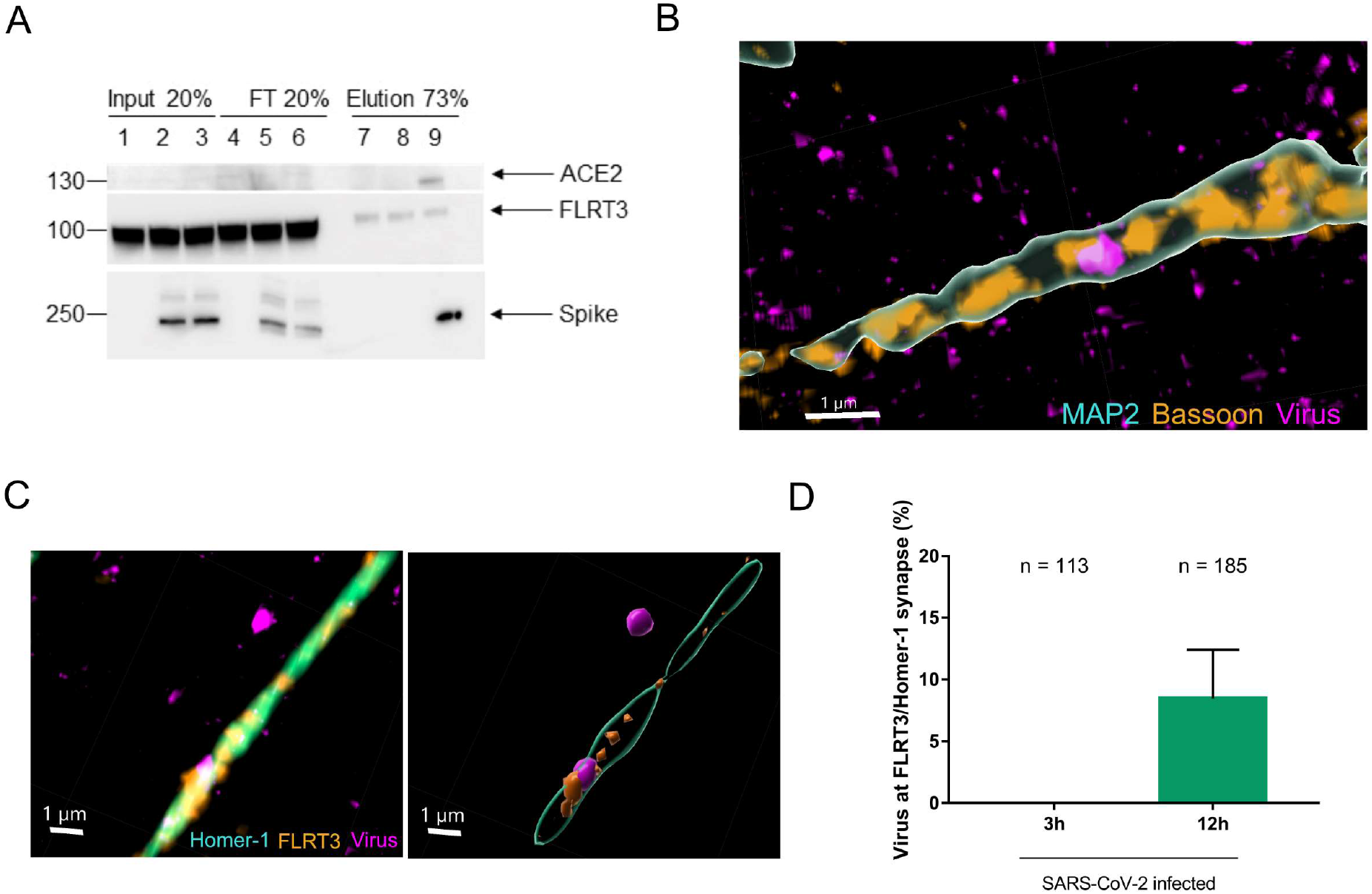
SARS-CoV-2 particles lodge at FLRT3-containing synapses. (**A**) Mature neurons derived from hNPCs were incubated with recombinant Spike protein for 3 h and immunoprecipitation of Spike was performed using protein A-coupled agarose beads. Western blotting for ACE2, FLRT3 and Spike shows that pull-down of the Spike protein was efficient, and that neuronal ACE2 could be retrieved associated to Spike, while undetectable from the input. However, no FLRT3 could be enriched upon Spike pull-down using this method. (**B-C**) Mature neurons derived from hNPCs were exposed to UV-inactivated SARS-CoV-2 N-mNeonGreen for 12 h, fixed, permeabilized and stained for the neuronal marker MAP2, the presynaptic proteins FLRT3 or Basson, and the post-synaptic marker Homer-1. Snapshots show viral particles associated to synaptic structures. (**D**) Quantification of the percentage of virus particles at FLRT3/Homer-1-positive synapses at 3 and 12 h post exposure. Data are mean +/- SD from 3 field of view. The n indicates the number of viral particles considered in each condition.

## Supplemental Tables

**Supplemental Table S1. Differential proteomics of single organoids.** The table shows fold-change (FC) values for each of the 3681 proteins identified and associated adjusted p values (pval) comparing four mock-infected organoids to four 1x SARS-CoV-2-infected organoids (1xCOV2) to three 10x SARS-CoV-2-infected organoids (10xCOV2).

**Supplemental Table S2. Post-translational modification analysis from single organoid proteomics.**

**Supplemental Table S3. Fold change of upregulated proteins from the synaptosome for each individual organoid**.

## Supplemental Movies

**Supplemental Movie S1. Live cell imaging of an active SARS-CoV-2 virus.** A SARS-CoV-2 N-mNeonGreen particle (Orange) was monitored on SynaptoRed-labeled neurons (Cyan) differentiated from hNPCs. Images were acquired every 5 s for 20 min using a spinning disk confocal microscope in a BSL-3 environment.

**Supplemental Movie S2. Live cell imaging of a dwelling SARS-CoV-2 virus.** A SARS-CoV-2 N-mNeonGreen particle (Orange) was monitored on SynaptoRed-labeled neurons (Cyan) differentiated from hNPCs. Images were acquired every 5 s for 20 min using a spinning disk confocal microscope in a BSL-3 environment.

## References

1. M. Gavriatopoulou et al., Organ-specific manifestations of COVID-19 infection. Clin Exp Med 20, 493–506 (2020).

2. S. Salinas, Y. Simonin, [Neurological damage linked to coronaviruses : SARS-CoV-2 and other human coronaviruses]. Med Sci (Paris) 36, 775–782 (2020).

3. I. J. Koralnik, K. L. Tyler, COVID-19: A Global Threat to the Nervous System. Ann Neurol 88, 1–11 (2020).

4. C. Iadecola, J. Anrather, H. Kamel, Effects of COVID-19 on the Nervous System. Cell 183, 16–27 e11 (2020).

5. J. Helms et al., Delirium and encephalopathy in severe COVID-19: a cohort analysis of ICU patients. Crit Care 24, 491 (2020).

6. A. Varatharaj et al., Neurological and neuropsychiatric complications of COVID-19 in 153 patients: a UK-wide surveillance study. Lancet Psychiatry 7, 875–882 (2020).

7. J. P. Rogers et al., Psychiatric and neuropsychiatric presentations associated with severe coronavirus infections: a systematic review and meta-analysis with comparison to the COVID-19 pandemic. Lancet Psychiatry 7, 611–627 (2020).

8. A. R. Antony, Z. Haneef, Systematic review of EEG findings in 617 patients diagnosed with COVID-19. Seizure 83, 234–241 (2020).

9. T. Kubota, P. K. Gajera, N. Kuroda, Meta-analysis of EEG findings in patients with COVID-19. Epilepsy Behav, 107682 (2020).

10. P. Nagu, A. Parashar, T. Behl, V. Mehta, CNS implications of COVID-19: a comprehensive review. Rev Neurosci 32, 219–234 (2021).

11. H. A. Baker, S. A. Safavynia, L. A. Evered, The ‘third wave’: impending cognitive and functional decline in COVID-19 survivors. Br J Anaesth 126, 44–47 (2021).

12. M. Taquet, J. R. Geddes, M. Husain, S. Luciano, P. J. Harrison, 6-month neurological and psychiatric outcomes in 236 379 survivors of COVID-19: a retrospective cohort study using electronic health records. Lancet Psychiatry 8, 416–427 (2021).

13. J. Hellmuth et al., Persistent COVID-19-associated neurocognitive symptoms in non-hospitalized patients. J Neurovirol, (2021).

14. G. Douaud et al., SARS-CoV-2 is associated with changes in brain structure in UK Biobank. medRxiv, (2022).

15. G. Blazhenets et al., Slow but Evident Recovery from Neocortical Dysfunction and Cognitive Impairment in a Series of Chronic COVID-19 Patients. J Nucl Med 62, 910–915 (2021).

16. A. C. Yang et al., Dysregulation of brain and choroid plexus cell types in severe COVID-19. Nature 595, 565–571 (2021).

17. A. Ramani, A. I. Pranty, J. Gopalakrishnan, Neurotropic Effects of SARS-CoV-2 Modeled by the Human Brain Organoids. Stem Cell Reports 16, 373–384 (2021).

18. E. Song et al., Neuroinvasion of SARS-CoV-2 in human and mouse brain. J Exp Med 218, (2021).

19. X. Qian, H. Song, G. L. Ming, Brain organoids: advances, applications and challenges. Development 146, (2019).

20. R. D. Hodge et al., Conserved cell types with divergent features in human versus mouse cortex. Nature 573, 61–68 (2019).

21. D. Chertow et al., SARS-CoV-2 infection and persistence throughout the human body and brain. Research Square, (2021).

22. X. Xie et al., An Infectious cDNA Clone of SARS-CoV-2. Cell Host Microbe 27, 841–848 e843 (2020).

23. A. Ramani et al., SARS-CoV-2 targets neurons of 3D human brain organoids. EMBO J 39, e106230 (2020).

24. M. Ferren et al., Hamster organotypic modeling of SARS-CoV-2 lung and brainstem infection. Nat Commun 12, 5809 (2021).

25. L. Bauer et al., The neuroinvasiveness, neurotropism, and neurovirulence of SARS-CoV-2. Trends Neurosci 45, 358–368 (2022).

26. M. Zivaljic et al., Poor sensitivity of iPSC-derived neural progenitors and glutamatergic neurons to SARS-CoV-2. BioRxiv, (2022).

27. F. Koopmans et al., SynGO: An Evidence-Based, Expert-Curated Knowledge Base for the Synapse. Neuron 103, 217–234 e214 (2019).

28. M. M. Silva et al., MicroRNA-186-5p controls GluA2 surface expression and synaptic scaling in hippocampal neurons. Proc Natl Acad Sci U S A 116, 5727–5736 (2019).

29. L. Lin et al., Electroencephalographic Abnormalities are Common in COVID-19 and are Associated with Outcomes. Ann Neurol 89, 872–883 (2021).

30. J. P. H. Burbach, D. H. Meijer, Latrophilin’s Social Protein Network. Front Neurosci 13, 643 (2019).

31. R. Sando, X. Jiang, T. C. Sudhof, Latrophilin GPCRs direct synapse specificity by coincident binding of FLRTs and teneurins. Science 363, (2019).

32. R. Sando, T. C. Sudhof, Latrophilin GPCR signaling mediates synapse formation. Elife 10, (2021).

33. M. L. O’Sullivan et al., FLRT proteins are endogenous latrophilin ligands and regulate excitatory synapse development. Neuron 73, 903–910 (2012).

34. D. G. May et al., A BioID-derived proximity interactome for SARS-CoV-2 proteins. bioRxiv, (2021).

35. W. Bakhache et al., Pharmacological perturbation of intracellular dynamics as a SARS-CoV-2 antiviral strategy. BioRxiv, (2021).

36. M. E. Coulter et al., The ESCRT-III Protein CHMP1A Mediates Secretion of Sonic Hedgehog on a Distinctive Subtype of Extracellular Vesicles. Cell Rep 24, 973–986 e978 (2018).

37. V. M. Corman et al., Detection of 2019 novel coronavirus (2019-nCoV) by real-time RT-PCR. Euro Surveill 25, (2020).

38. D. Bouyssie et al., Proline: an efficient and user-friendly software suite for large-scale proteomics. Bioinformatics 36, 3148–3155 (2020).

39. S. Wieczorek, F. Combes, H. Borges, T. Burger, Protein-Level Statistical Analysis of Quantitative Label-Free Proteomics Data with ProStaR. Methods Mol Biol 1959, 225–246 (2019).

40. Y. Perez-Riverol et al., The PRIDE database resources in 2022: a hub for mass spectrometry-based proteomics evidences. Nucleic Acids Res 50, D543–D552 (2022).

41. N. Hulstaert et al., ThermoRawFileParser: Modular, Scalable, and Cross-Platform RAW File Conversion. J Proteome Res 19, 537–542 (2020).

42. S. Degroeve et al., ionbot: a novel, innovative and sensitive machine learning approach to LC-MS/MS peptide identification. BioRxiv, (2021).

